# Novel missense mutation E585K in retinitis pigmentosa leads to compromised *RPGR* splicing diversity

**DOI:** 10.1101/2020.05.22.109884

**Authors:** Yan-Shan Liu, Ji-Feng Wan, Chun-Yan Ren, Zhou-Heng Xu, Xu-Bin Pan, Jia-Qi Pan, Ruo-Nan Gao, Shao-Qiang Liu, Jia-Li Zhang, Qian-Hao Yao, Ji-Hong Wang, En-Min Li, Jun-Hua Rao, Ping Hou, Jian-Huan Chen

## Abstract

Mutations in the retinitis pigmentosa GTPase regulator (*RPGR*) gene, are the major cause of X-linked retinitis pigmentosa (RP). Herein we used whole-exome sequencing to screen possible novel *RPGR* mutations in RP patients, and identified a novel missense mutation E585K in a patient with early onset but slow disease progression, and a frameshift deletion E998Gfs*78 in a patient with RP sine pigmento and high myopia. Intriguingly, bioinformatic analysis indicated that E585K probably affected *RPGR* RNA splicing instead of the protein sequence directly. Mini-gene assays in 293T cells revealed that splicing events of the E585K mutant were found to be also exist in wildtype, but with a shifted pattern. In the E585K mini-gene usage of an upstream alternative 5′ splice site (5′ ss) of exon 14 was enhanced, and other splicing events were suppressed, including the canonical 5′ ss of exon 14, skipping of exon 14/15 and retention of intron 14. As a result, *RPGR* splicing products of the E585K mini-gene were predominated by transcripts containing a 4-bp deletion, with a small fraction of in-frame transcripts containing a retended intron 14, which might explain the slow disease progression in the patient carrying the mutation. RNA-Seq analysis further confirmed existence of these splicing events in endogenous *RPGR* RNA in human retina, pointing to compromised splicing diversity in the E585K mutant. Our findings thus added to the understanding of genotype-phenotype correlation in RP, and suggested that compromised *RPGR* splicing diversity might play a role in molecular mechanism of the disease.

## Introduction

Retinitis pigmentosa (RP, MIM 268000) is a group of inherited retinal degenerative diseases characterized by night blindness due to the progressive loss of rod photoreceptors, followed by complete blindness due to the further loss of cones (Hamel 2006, Tsang and Sharma 2018). X linked retinitis pigmentosa (XLRP) represents the most severe and prevalent form owing to its early onset and severity of progress (Tee et al. 2018, Tsang and Sharma 2018). To date, six loci have been associated with XLRP, yet only three genes have been identified (Lyraki et al. 2016, Ferrante et al. 2001, Schwahn et al. 1998, Meindl et al. 1996), among which the retinitis pigmentosa GTPase regulator (*RPGR*) gene [MIM 312610] accounts for 70-90% of XLRP and 15-20% of simplex RP cases, making it one of the most common causes of RP (Parmeggiani et al. 2017, Khanna 2018).

*RPGR* locates at Xp11.4 and undergoes complex alternative splicing in its RNA expression. Two major transcripts, the constitutive *RPGR*_ex1-19_ and *RPGR*_ORF15_, have been extensively studied in XLRP (Megaw et al. 2015a, Charng et al. 2019). *RPGR*_ORF15_ shares the same exon 1-14 with *RPGR*_ex1-19_ and uses an alternative large ORF15 exon. *RPGR*_ORF15_ encodes a purine-rich, highly repetitive sequence followed by a basic C terminal domain with unknown function (Patnaik et al. 2015). The *RPGR* ORF15 region is also a mutation hotspot, accounting for around 60% of *RPGR* pathogenic mutations (Beltran et al. 2014, Vervoort et al. 2000). Most of the mutations in this region are one to two, or four to five bp deletions in exon, resulting in a truncated protein with the loss of C terminal domain (Cehajic Kapetanovic et al. 2019). The Constitutive *RPGR*_ex1-19_ is widely expressed while *RPGR*_ORF15_ is highly expressed in retina. Overexpression of *RPGR*_ex1-19_ results in severe retinal degeneration, suggesting a delicate ratio between two isoforms, regulated by alternative splicing, is important for integrity and function of retina (Liu and Zack 2013, Wright et al. 2011).

In the current study, our whole-exome sequencing (WES) in male RP patients identified two mutations in *RPGR* (NM_001034853), including a novel missense mutation c.1753G>A (E585K) in a patient with early onset but slow disease progression, and a frameshift deletion E998Gfs*78 (Vervoort et al. 2000) in a patient with RP sine pigmento and high myopia. The E585K mutant mini-gene showed compromised *RPGR* splicing diversity, resulting in predominant frameshift transcripts and a small fraction of transcripts with intron 14 retended. It was noteworthy that all of these splicing variants were also found in endogenous *RPGR* expression across multiple human cell types, but the compositions of splicing variants were altered compared to wildtype. Our findings indicated that compromised splicing diversity instead of biogenesis of new disease-causing transcripts in *RPGR* expression could play a potential role in molecular mechanism of RP.

## Methods

### Clinical examination and sample collection

This study was approved by the Ethics Committee of Jiangnan University, and was conducted in accordance with the Declaration of Helsinki. Written consent was obtained from all participants after explanation of the nature of the study. Eleven male unrelated Chinese nonsyndromic RP patients were included in the current study after comprehensive ophthalmic examination. Peripheral blood was collected from all of the patients, and genomic DNA was extracted using TIANamp Blood DNA Maxi Kit (TIANGEN, Beijing, China).

### Whole exome sequencing

Genomic DNA (3 μg per individual) from selected individuals was taken for WES by Novogene (Tianjin, China). The whole exome was captured by an Agilent SureSelect All Human exon v5.0 kit (Agilent, Santa Clara, US.), and sequenced on an Illumina Hiseq X-10 (Illumina, Hayward, US.) with a paired-end 150 bp length configuration.

### Analysis of whole exome sequencing data

The WES data were analyzed as previously described (Chen et al. 2016). Quality of raw sequencing data was analyzed by using FastQC (https://www.bioinformatics.babraham.ac.uk/projects/fastqc/). BWA (Burrows-Wheeler Aligner, Version 0.7.6) was used for mapping short reads against the Human Reference Genome hg19 from UCSC Genome Browser (http://genome.ucsc.edu/)(Li and Durbin 2009). Variants were called and recalibrated according to the best practice pipeline by GATK (version 3.6), and then annotated using ANNOVAR (Wang et al. 2010). Minor allele frequencies (MAFs) of genetic variants were estimated using available data from dbSNP138 (http://www.ncbi.nlm.nih.gov/snp/), the 1000 Genomes Project (http://www.1000genomes.org), the NHLBI Exome Sequencing Project (ESP, http://evs.gs.washington.edu/EVS/), the exome array design data (http://genome.sph.umich.edu/wiki/Exome_Chip_Design) (Huyghe et al. 2013), and the Exome Aggregation Consortium (ExAC, http://exac.broadinstitute.org/) and gnomAD (https://gnomad.broadinstitute.org/) data. SIFT (http://genetics.bwh.harvard.edu/pph/) (Kumar et al. 2009) and PolyPhen (http://sift.jcvi.org/www/SIFT_chr_coords_indels_submit.html) (Ramensky et al. 2002) programs were used to predict the functional impact of remaining variants on the encoded protein. The WES detected variants were then screened for possible mutations in known genes resulting in retinitis pigmentosa according to Retinal Information Network (RetNet, https://sph.uth.edu/retnet/).

### Sanger sequencing

The sequences of genes with selected variants were obtained from the NCBI reference sequence database (http://www.ncbi.nlm.nih.gov/Refseq). Primers were then designed to confirm selected variants; detailed information of primers was shown on Table 1. Primers were designed accordingly by Primer 3 are summarized in **Table S1**. Genomic DNA from patients was amplified by Polymerase chain reaction (PCR) using PrimeSTAR HS Premix (Takara Bio) on a Mastercycler nexus (Eppendorf, Hamburg, Germany) as described previously (Chen et al. 2016). Sequence alignment and analysis of variations were performed using the NovoSNP program (Weckx et al. 2005).

**Table 1.** Characteristics of *RPGR* mutations resulting from filtering of exome sequencing data.

NF, not found; NA, not available
| No. | Chromosomal position<sup>a</sup> | Gene symbol | mRNA | cDNA change<br>(Protein change) | Functional prediction |  | dbSNP | Public /Control exomes | Sanger Sequencing |
| --- | --- | --- | --- | --- | --- | --- | --- | --- | --- |
|  |  |  |  |  | SIFT | Polyphen |  |  |  |
| 1 | chrX:38147114 | *RPGR* | NM_001034853 | c.1753G>A<sup>d</sup><br>(E585K) | tolerated | Benign | NF | NF | Confirmed |
| 2 | chrX:38145255 | *RPGR* | NM_001034853 | c.2993_2997delAAGGG<br>(E998Gfs*78) | NA | NA | NF | NF | Confirmed |

<sup>a</sup>Chromosomal positions are given according to the UCSC hg19 reference assembly.
<sup>b</sup>The mutation is at the last base of exon 14, 1 bp upstream of the canonical 5'ss.

### In-house control exomes

A dataset of 150 in-house control exomes, which was consolidated from multiple previous WES studies performed by our laboratory, was used to filter out variants that were not related to the disease phenotype. all control individuals had no sign or family history of retinitis pigmentosa and other inherited retinal diseases.

### Mini-gene plasmid construction

To find out whether the presumed missense mutation affected *RPGR* splicing, a *RPGR* mini-gene was constructed based on pEGFP-N3 (TAKARA Bio, Kusatsu, Japan). Genomic sequence spanning from exon13 to exon 15, including intron 13 and 14 of *RPGR* containing the wildtype and mutant of c.1753G>A were amplified from genomic DNA of an unrelated healthy control and patient 1 respectively, and inserted together with a Kozak sequence into the N terminus of EGFP. The *RPGR* mini-genes were expected to generate a fusion protein with EGFP.

### Cell culture and transfection

Human 293T cells and ARPE-19 cells (American Tissue Culture Collection, Manassas, US.) were cultured in 6-well plates (Corning, Corning, US.) in Dulbecco’s modified Eagle’s medium: Ham’s F-12 (DMEM/F12, ThermoFisher Scienticfic, Waltham, US.) supplemented with 10% fetal bovine serum (FBS, ThermoFisher Scienticfic, Waltham, US.) and antibiotics (100 units/ml penicillin G and 100 μg/ml streptomycin sulfate, ThermoFisher Scienticfic, Waltham, US.) at 37°C in a 5% CO_2_ incubator. 293T cells were transfected with 2.5 μg of wild-type or mutant mini-gene plasmids using Lipofectamine 3000 (ThermoFisher Scienticfic, Waltham, USA) following the manufacturer’s instruction.

### Reverse transcription and PCR

Forty-eight hours after transfection total RNA was extracted using RNApuro Tissue & Cell kit (CWBIO, Beijing, China), and reverse transcription (RT) was performed using PrimeScript RT reagent Kit with gDNA Eraser (TAKARA Bio, Kusatsu, Japan). Primers were designed to specifically amplify cDNA from transfected vectors and endogenous *RPGR* from 293T and ARPE-19 cells respectively. The sequences of PCR products were confirmed using Sanger sequencing. Primers for expression analysis are listed in **Table S1**.

### Immunoblotting

Cells were lysed and total protein was extracted using protein lysis buffer (Biyotime, Shanghai, China) supplemented with a protease inhibitor (CWBIO, Beijing, China). Protein samples were run on a 12% acrylamide gel, and immunoblotting was performed using antibodies described above. Antibodies used in this study include mouse-anti-EGFP (AE012, ABClonal, Wuhan, China); HRP-conjugated mouse monoclonal anti-GAPDH (KC-5G5, Kangchen, Shanghai, China).

### Splicing prediction

Splicing prediction of wild type and mutant variants was conducted using Human Splicing Finder (HSF, http://www.umd.be/HSF/) (Desmet et al. 2009).

### Analysis of published retina RNA-Seq data

We used published RNA-seq data to investigate of *in vivo RPGR* splicing (Ratnapriya et al. 2019). Original data was downloaded from Gene Expression Omnibus (SRA, www.ncbi.nlm.nih.gov/geo) under access No. GSE115828. Original RNA-seq reads were aligned to human whole genome (hg38) with STAR (v2.7.0) (Dobin and Gingeras 2015), and visualized with IGV (v 2.4.17) (Robinson et al. 2011).

## Results

### WES identified two RPGR mutations

After analysis of exome sequencing data from the eleven male RP patients, we found two *RPGR* mutations in two RP patients. The summary of original exome sequencing data and detected variants of the two RP patients with *RPGR* mutations were shown in **Table S2** and **S3**. The two hemizygotic *RPGR* mutations in two unrelated male Chinese RP patients, after exclusion of any possible mutation in other known RP causative genes **(Figure 1A**). Patient 1 had a novel missense mutation, c.1753G>A (E585K) and Patient 2 had a reported five-bp frameshift deletion, c.2993_2997del (E998Gfs*78) (Vervoort et al. 2000). The novel missense mutation was not found in either the public databases listed in the methods, or our control exomes. Sanger sequencing further confirmed both mutations in these two patients (**Figure 1B** **and** **1C**). Multiple sequence alignment showed that for E585K, the mutated glutamine residue at codon 585 was not evolutionarily conserved (**Figure 1D**), and SIFT and Polyphen predicted the mutation to be benign (**Table 1**). In addition, the mutated nucleotide at 1753 position of the cDNA was not conserved among mammals, either (**Figure 1E**). However, c.1753G>A was at the last base of exon 14, one bp upstream to the canonical splice donor (5′ss), and thus might had potential impact on RNA splicing.

**Figure 1.**
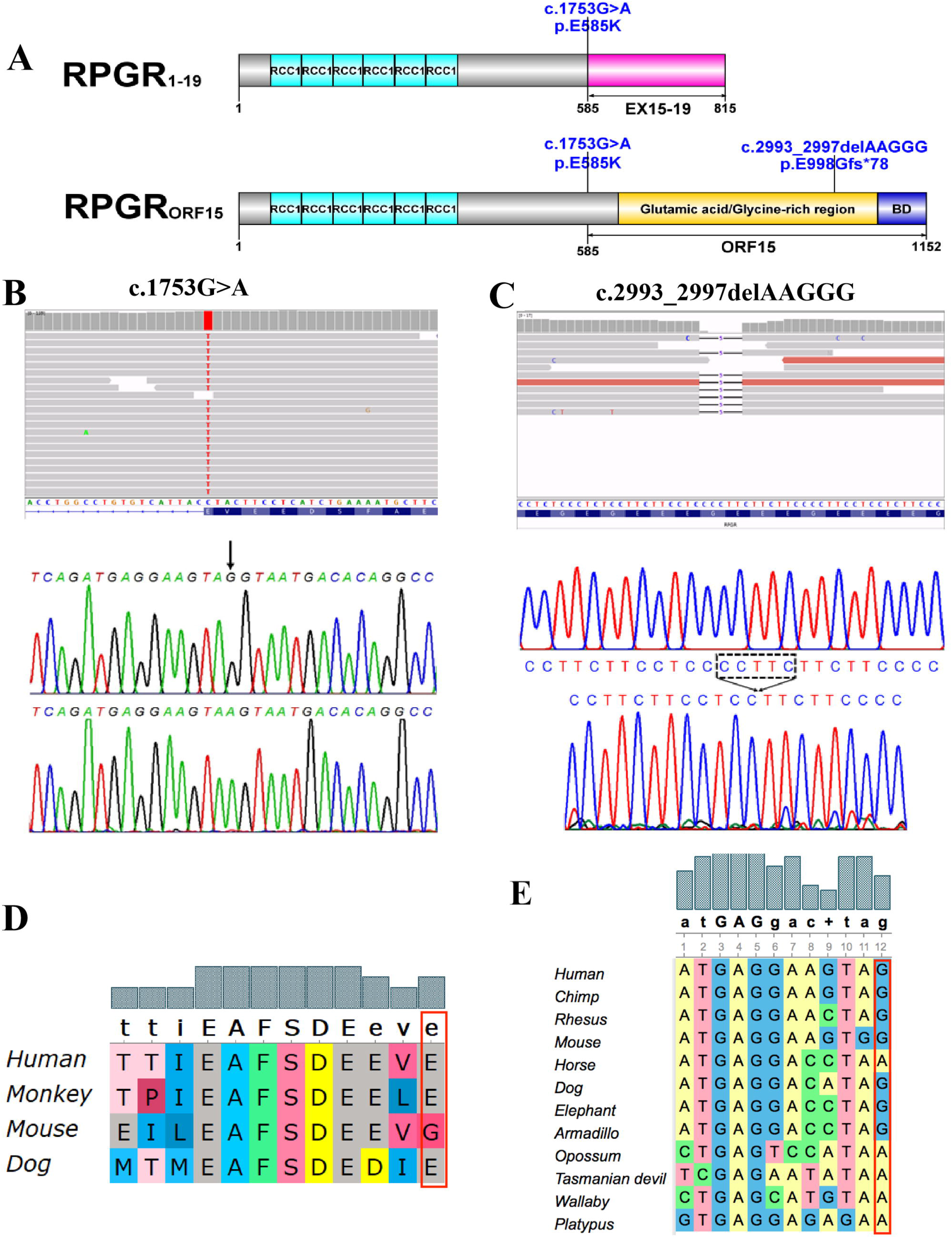
Two novel *RPGR* mutations identified by WES in RP patients. (A). Location of the two mutations on RPGR_1-19_ and/or RPGR_ORF15_. The Regulator of Chromosome Condensation (RCC1) like domains on both isoforms, the Glutamic acid/Glycine-rich domain and Basic domain (BD) are shown as well. (B) and (C). Alignment of exome data followed by Sanger sequencing reveals two novel *RPGR* mutations in two male Chinese RP patients. (D). Alignment of amino acids reveals low conservation of the glutamate residue mutated in E585K among mammals. (E) Alignment of genomic sequence shows the last base of human *RPGR* exon 14 is not conserved among mammals.

### Clinical phenotyping results of the RP patients with RPGR mutations

Clinical phenotyping analysis was then conducted in the two RP patients with *RPGR* mutations (**Table 2**). Patient 1 with the E585K missense mutation was a 50-year-old male, with self-reported bilateral night blindness and blurry vision that started at around 10 years old. His symptoms progressed slowly after disease-onset, but deteriorated significantly over the past ten years. He had no other genetic eye disease or syndrome except posterior subcapsular cataracts, with normal intraocular pressure (< 21 mmHg). Fundus examination revealed bilateral severe, diffuse, granular appearance and atrophy of retinal pigment epithelium (RPE), and bone-spicule pigmentation, as well as arteriolar attenuation in mid-peripheral retina. Furthermore, optical coherence tomography (OCT) showed mild thinning of inner retinal layer and severe thinning of outer retinal layer (**Figure 2A** **and** **2B**).

**Table 2.** Clinical features of RP patients with *RPGR* mutations in the current study.

| Patient<br>(Mutation) | Sex | Age at<br>recruitment | Age<br>of<br>onset | Visual<br>acuity |  | Details of RP phenotype | Other ocular<br>symptoms |
| --- | --- | --- | --- | --- | --- | --- | --- |
|  |  |  |  | OD | OS |  |  |
| Patient 1<br>(c.1753G>A<sup>a</sup>) | Male | 50 | 10 | LP | LP | Bilateral; night blindness; severe, diffuse, granular appearance and atrophy of RPE; bone-spicule pigmentation; arteriolar attenuation in mid-peripheral retina | Posterior subcapsular cataracts |
| Patient 2<br>(c.2993_2997del) | Male | 20 | 7-8 | HM | 0.02 | Bilateral; night blindness; mild bone-spicule pigmentation, and unrecordable ERG | High myopia |

NA: not available
HM, hand motion; LP, light perception
RPE, retinal pigment epithelium; ERG, electroretinogram
<sup>a</sup>This variant is at the last base of exon 14, 1bp upstream to the donor site, and thus might had potential impact on RNA splicing.

**Figure 2.**
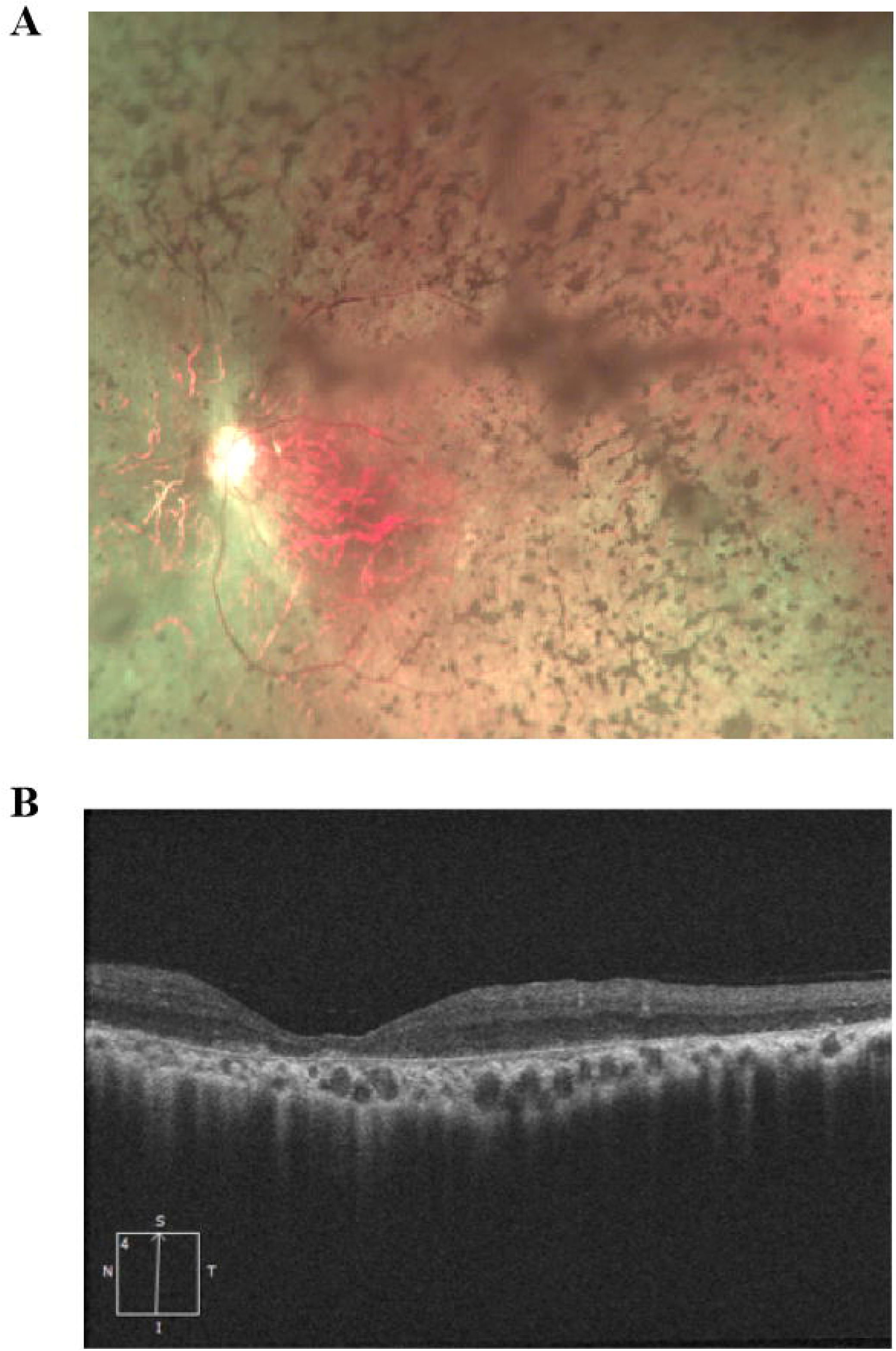
Clinical examination of patient 1 with the *RPGR* c.1753G>A mutation. (A) Fundus photos of Patient 1, showing bone-spicule pigmentation. Moderate senile cataract is also observed in the patient. (B) Optical coherence tomography of patient 1 shows mild inner retinal layer thinning and severe outer retinal layer thinning.

Patient 2 was a 20-year-old male proband (III-3) from a three-generation XLRP family (**Figure S1A**), in which his younger brother (III-4) and one of his cousins (I-1) were also affected by RP. He suffered from gradually decreased visual acuity starting at six years old, with night blindness started from seven to eight years old. Fundus examination showed that he had bilateral unrecordable electroretinogram (ERG), but lacked characteristic bone spicule pigmentation, and thus was diagnosed as RP sine pigmento (**Figure S1B and S1C)**. No other ocular disease or syndrome was found except high myopia.

### Splicing analysis revealed splicing diversity near RPGR exon 14

To better understand the splicing of near the *RPGR* Exon 14, we then checked *RPGR* mRNA expression in human 293T cells and ARPE-19 cells (**Figure 3A**). Our Sanger sequencing results indicated that in both cell lines, there were probably two distinct 5′ss in (**Figure 3B)**. In addition to a canonical 5′ss at the boundary of exon 14 (E14c), a novel alternative 5′ss was found at 4 bp upstream of it. As a result, splicing led to a 4 base frameshift deletion and a shorter exon 14 (E14s). Intriguingly, it was noted that the relative level of the alternative 5′ss usage and E14s was lower in ARPE-19 cells compared to 293T cells, probably in line with a higher expression level of RPGR_ORF15_ in the retina. Furthermore, by using published human retina RNA-seq data, we confirmed that retina expressed transcripts that lacked the last 4 bp of exon 14 (**Figure S2**), which further supported the existence of the intrinsic alternative 5′ss and E14s in normal human retina. In addition, RT-PCR followed by Sanger sequencing and RNA-seq also showed that, skipping of exon 14 and 15, and retention of intron 14 could also be observed in endogenous *RPGR* RNA in these human cells (**Figure 3C, 3D**, **and S3**). Taken together, these findings demonstrated high levels of splicing diversity near exon 14 in endogenous *RPGR* expression.

**Figure 3.**
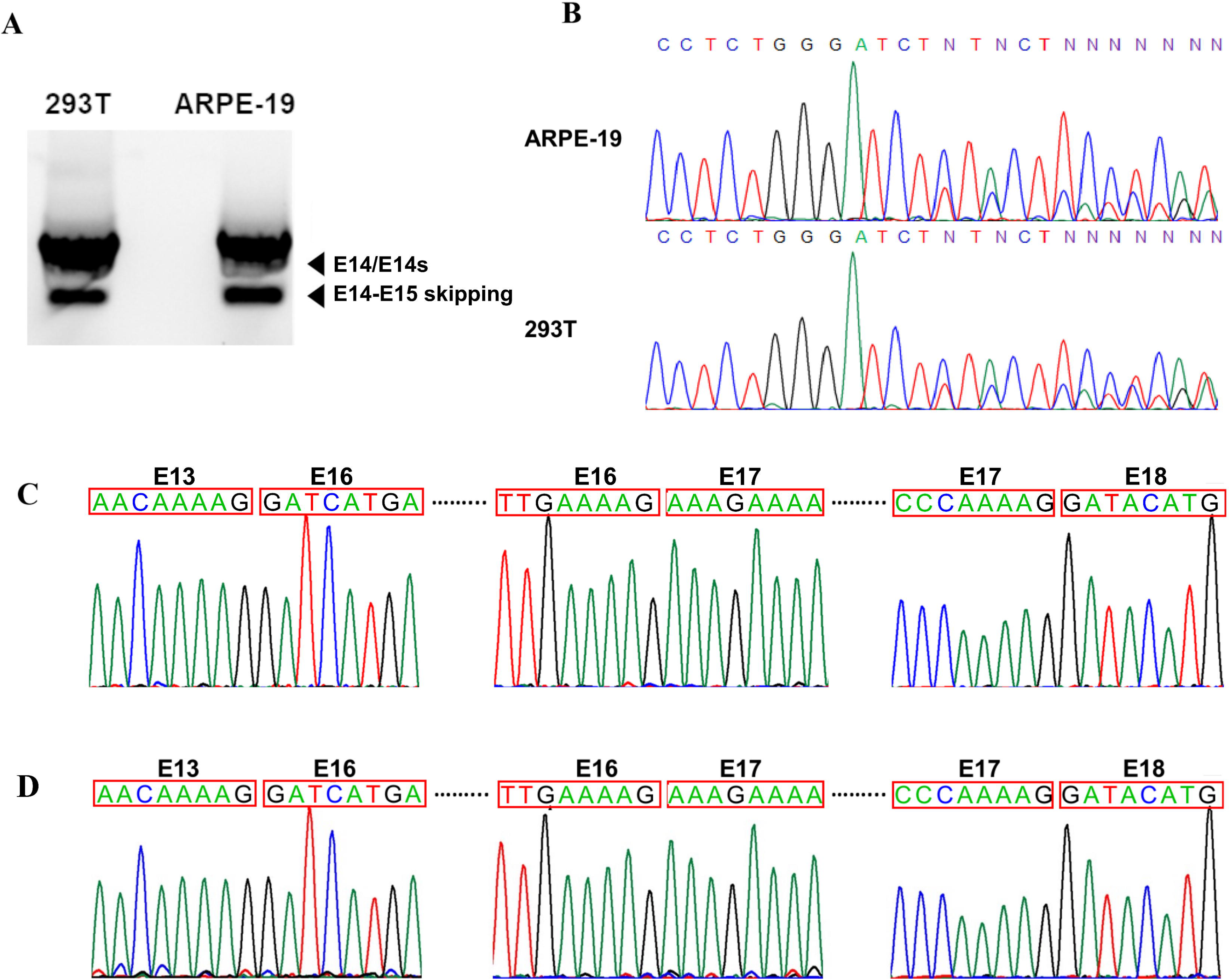
Existing of the alternative exon 14 5′ss and exon 14 skipping in endogenous *RPGR* RNA in ARPE-19 and 293T cells. (A) RT-PCR of ARPE-19 and 293T cells shows multiple endogenous *RPGR* splicing events. (B) Sanger sequence of endogenous *RPGR* RNA in ARPE-19 and 293T cells shows existence of both the canonical and the novel alternative 5′ss in *RPGR* E14. Sanger sequencing confirmed skipping of exon 14 and 15 in endogenous *RPGR* RNA in ARPE-19 (C) and 293T cells (D).

### The E585K mutation led to compromised RPGR splicing diversity

The missense mutation E585K occurs one bp upstream to the canonical splice 5′ss of exon 14. Therefore, the E585K mutation was selected for further functional analysis to explore its potential impact on RNA splicing. To analyze whether and how this mutation disrupted splicing of *RPGR*, the entire genomic region spanning *RPGR* Exon 13, intron 13, exon 14, intron 14, and exon 15 were cloned into pEGFP-N3 (**Figure 4A**), and overexpressed in human 293T cells. RT-PCR showed different splicing patterns between wildtype and the c.1753G>A mutant **(Figure 4B)**. Similar to endogenous *RPGR* RNA, the wildtype mini-gene was found to exhibit high splicing diversity with four distinct splicing events (**Figure 4B-E**). As confirmed by Sanger sequencing, the exon 14 skipping, E14c/E14s, and intron 14 retention accounted for 34.9%, 34.5% and 30.5% of the total wildtype splicing products respectively, while in the E585K mini-gene, E14s became predominant (86.9%) with a small fraction of transcripts containing the retended intron 14 (13.1%) (**Figure 4C, 4E-G**). Western-blot indicated that fusion in-frame EGFP protein was detected for wildtype mini-gene but not for the E585K mutant (**Figure 4D**). Human splicing finder predicted that the mutation a novel binding site for SRp55 protein (**Table S4**), which might contribute to compromised splicing diversity and enhanced usage of the alternative 5′ss.

**Figure 4.**
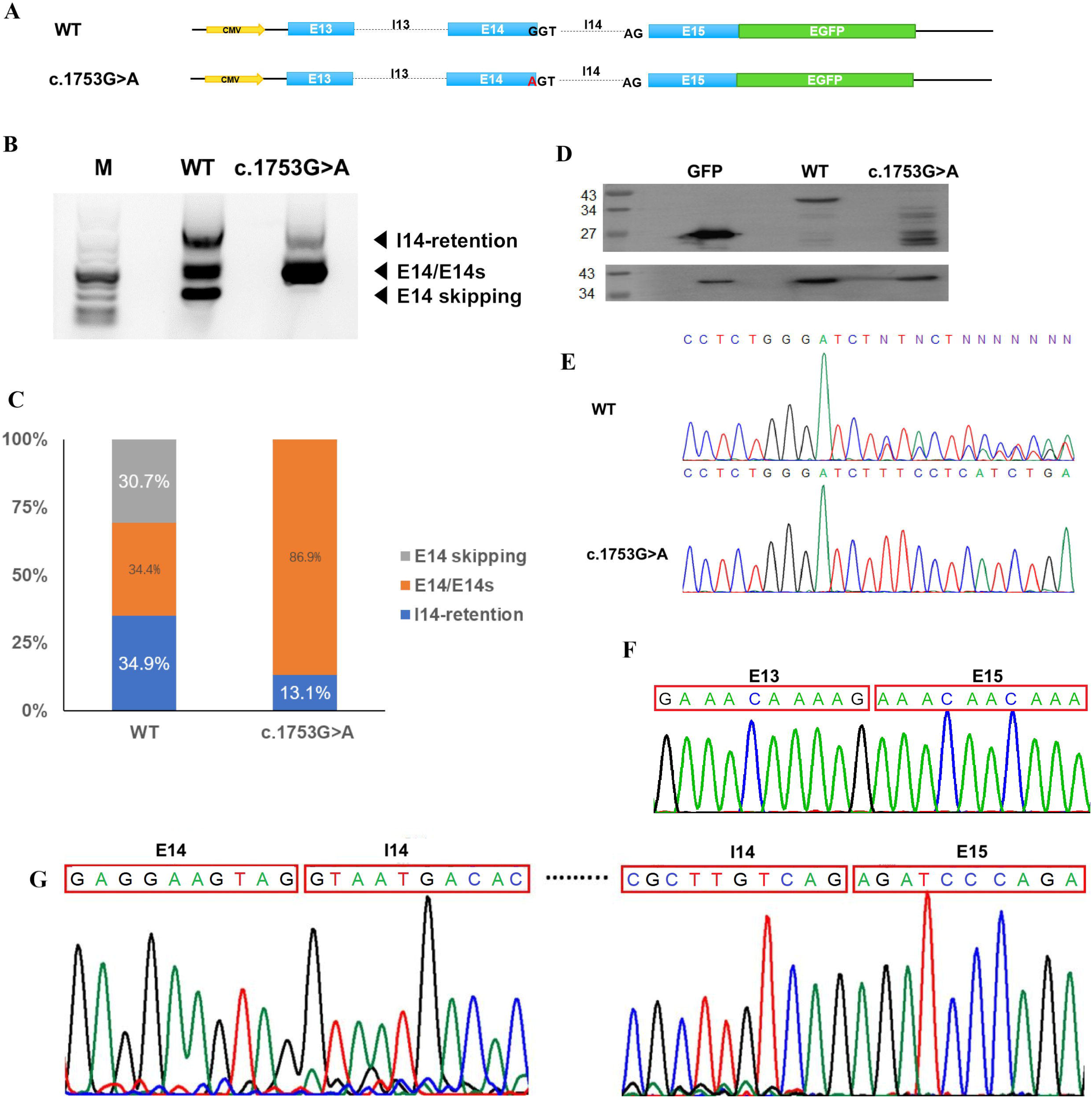
Compromised *RPGR* splicing diversity in the c.1753G>A mutant. (A) Construction of mini-gene to investigate the effects of c.1753G>A on splicing. (B) RT-PCR of 293T cells transfected with wildtype or c.1753G>A mini-genes show distinct *RPGR* splicing patterns. (C) Quantification of the RT-PCR results shown in (B). (D) Western blot of 293T cells transfected with EGFP, wildtype or c.1753G>A mini-gene. The c.1753G>A mutation suppresses the canonical 5′ss in E14, resulting in undetectable fusion GFP protein with canonical RPGR. (E) Sanger sequencing confirms existence of both the canonical and the novel alternative 5′ss in *RPGR* exon 14 in wildtype. (F) Sanger sequencing confirms skipping of exon 14 in wildtype, which is suppressed in the c.1753G>A mutant. (G)Sanger sequencing confirms retention of intron 14 in both wildtype and the c.1753G>A mutant.

## Discussion

In the current study, our WES identified two hemizygotic *RPGR* mutation in two male RP patients, including a novel E585K missense mutation in a patient with early onset but slow disease progression, and a frameshift deletion E998Gfs*78 in a patient with RP sine pigmento. Functional studies showed the missense mutation E585K suppressed the canonical 5′ss and skipping of E14, and enhanced an alternative 5′ss, leading to compromised splicing diversity and predomination of E14s-containing transcripts. Our results thus added to *RPGR* mutation spectrum and genotype-phenotype correlation in RP, and moreover, implicated the compromised *RPGR* splicing diversity could play a potential role in in RP etiology.

*RPGR* undergoes complex alternative splicing in its RNA expression, and gives rise to multiple isoforms (Charng et al. 2019). Both RPGR_ex1-19_ and RPGR_ORF15_ isoforms localize to the connecting cilium of photoreceptor, and play a role of sorting and trafficking proteins from inner to outer segment (Zhang et al. 2019). However, emerging evidence indicates that RPGR_ORF15_ is more closely involved in RP (Talib et al.). RPGR_ORF15_ is highly expressed in retina, and is formed by exon 15 extending into intron 15 due to skipping of splicing, generating a repetitive sequence rich in Glutamic acid and Glycine residues and a highly conserved basic C terminus (Tee et al. 2016). Previous studies have shown that the domain interacts with WHRLN, a scaffold protein that plays a role in assembling of cytoskeleton in ear and photoreceptor (Zhang et al. 2019) (Wright et al. 2012). By far, all disease causing mutations are found to affect RPGR_ORF15_, which is highly expressed in retina (Megaw et al. 2015b). Previous studies have shown that most of *RPGR* mutations result in truncated RPGR_ORF15_ and loss of C-terminal domain. In line with these previous findings, the two *RPGR* mutations identified in our RP patients also affected RPGR_ORF15_ function. As shown in **Figure 1A**, the frameshift deletion c.2993_2997del (p.E998Gfs*78) mutation in exon 15 led to a truncated glutamic acid/Glycine-rich region and loss of the basic domain, which resulted in RP sine pigmento with high myopia in Patient 2. Before our study, there has be no RPGR mutation to be associated with RP sine pigmento. In addition, the presumed missense mutation E585K probably disrupted the splicing of normal ORF15 exon, and caused typical RP phenotypes in Patient 1. Such findings therefore added to the mutation spectrum affecting the ORF15 exon, and further underlined importance of the alternative exon in RP.

Alternatively splicing plays a role in cell differentiation, lineage determination, tissue identity acquisition and maintenance, and organ development (Baralle and Giudice 2017). A delicate splicing needs spliceosome and splicing cis-elements, including 5′ss, branch point sequence, polypyrimidine tract and splice acceptors (Ramanouskaya and Grinev 2017). Alternations of these elements could affect the efficiency, or disrupt proper splicing. In the current study, we confirmed a novel alternative 5′ss by both RT-PCR and RNA-Seq, which resulted in a 4-bp frameshift deletion and a shorter exon 14 (E14s). Interestingly, it is noted that E14s existed in multiple cell types, with a high abundance in human 293T cells (**Figure 3B**). In contrast, in RPE cells where RPGR_ORF15_ was highly expressed, the abundance of E14s was much lower, indicating the potential importance of *RPGR* splicing regulation near exon 14.

The consensus sequence of 5′ss is CAG/GUAAGUAU, in which the GU after CAG is the donor site (Ohno et al. 2018). The last base before donor site is conserved as ‘G’ in 80% of 5′ss (Demirci et al. 2004). In the present study, the c.1753G>A (E585K) mutation, which was predicted to introduce a missense mutation at protein level, located one bp upstream of the canonical 5′ss of exon 14, and thus might affect splicing. Our *in vitro* mini-gene assay confirmed such possibility by showing the first evidence of a presumed benign ‘missense’ mutation in exon 14 that in fact could disrupt *RPGR* function. Previous studies showed that mutations at the last base in exons usually result in exon skipping or intron retention. For example, exon skipping could be found in around 70% mutations at the last base of exons in cancers (Jung et al. 2015). In RP, a mutation c.213G>A (G52R) has been reported to disrupt *RPGR* splicing by generating an aberrant transcript due to skipping of exon 2 (Demirci et al. 2004). However, it didn’t fit the whole story in our study. Our mini-gene assay showed that wildtype *RPGR* has a complex pattern near exon 14 that give rise to multiple splicing products in this region, including skipping of exon 14, retention of intron 14, splicing with the canonical 5′ss, as well as with the alternative 5′ss. In RPE, splicing with the canonical 5′ss showed a high priority, which generates the ‘canonical’ functional *RPGR*. Splicing with alternative 5′ss, which might generate frameshift *RPGR* transcripts, were found to exist in multiple human cell types. These findings suggest that both skipping of exon 14 and the alternative 5′ss exist in wildtype and various normal human tissues or cells, and are unlikely to cause RP by the two events directly. Instead, E585K enhanced the alternative 5′ss, and suppressed all the other events. As a result, it led to predominant transcripts containing E14s, and thus compromised splicing diversity (**Figure 5**). Such findings together with point to the importance of delicate balance in alternative splicing regulation, which maintain *RPGR* function.

**Figure 5.**
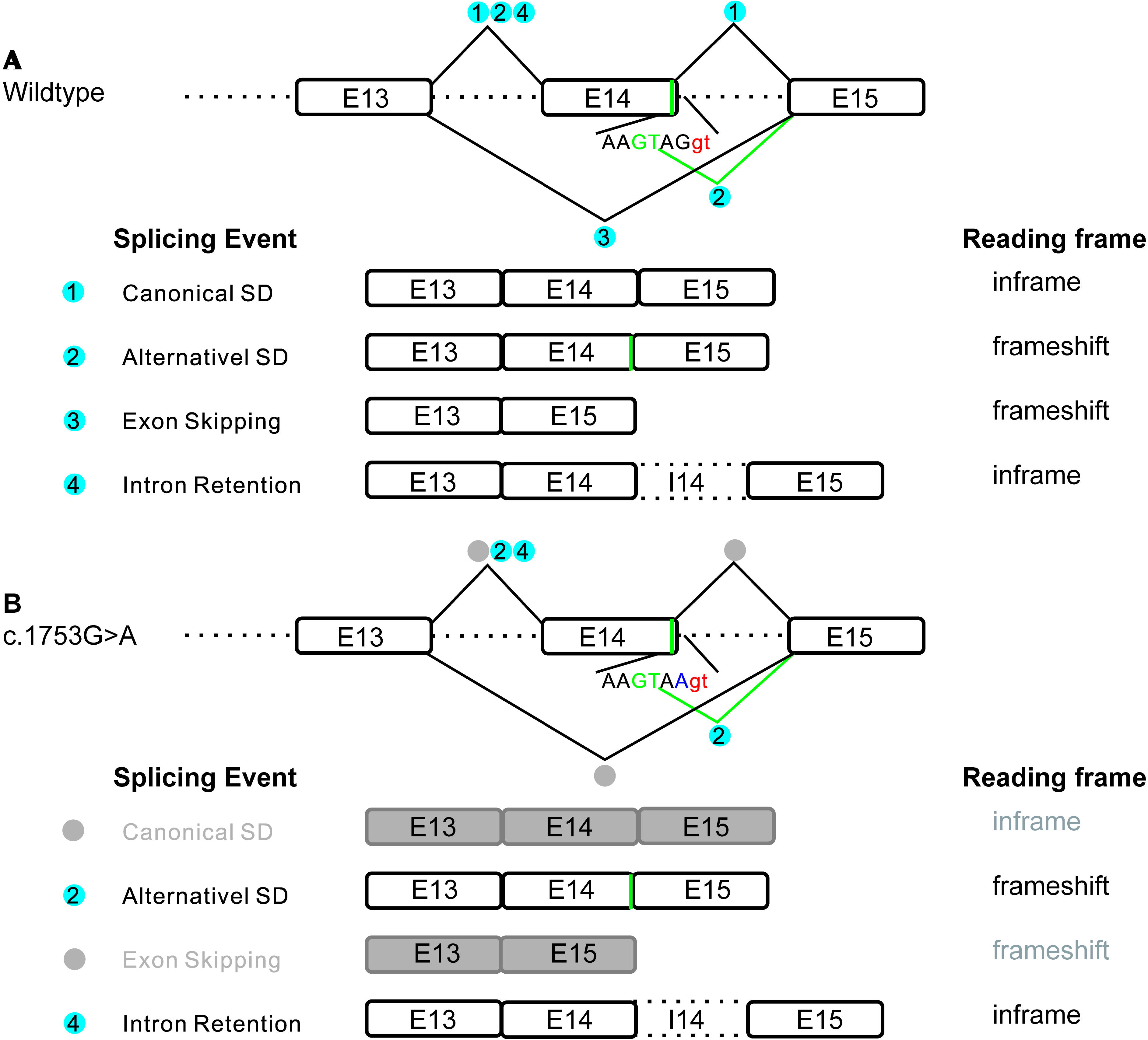
A schematic diagram illustrating compromised *RPGR* splicing diversity due to the c.1753G>A mutation. (A) For the wildtype *RPGR*, the canonical 5′ss (red) generates a canonical *RPGR* with exon 13, 14 and 15. The alternative 5′ss (green) generates a transcript with the last 4bp of exon 14 deleted (E14s), resulting in a frameshift *RPGR*. Skipping of exon14 generates a frameshift transcript, resulting in nonsense mediated mRNA decay. And retention of intron 14 produces an in-frame but longer transcript. (B) Splicing events of the canonical 5′ss and skipping of exon14 are suppressed due to the c.1753G>A mutation (purple), giving rise to predominant transcripts containing E14s with a small fraction of transcripts containing the retended intron 14, and thus compromised splicing diversity.

In summary, we identified two mutations in *RPGR* in RP patients with different clinical features, among which a novel missense mutation c.1753G>A (E585K) was found to cause compromised splicing diversity of *RPGR* exon 13-15 junctions in the ORF15 region, and led to predomination of alternative spliced transcripts. Our findings expand our knowledge about the mutation spectrum in RP, and provide us a new insight into alternative splicing regulation of *RPGR* in RP.

## Acknowledgements

The authors thank the patients and family members for collaborating with this study. This study was supported in part by grants from the National Natural Science Foundation of China (No. 31671311 and 81371033), the National first-class discipline program of Light Industry Technology and Engineering (LITE2018-14), the “Six Talent Peak” Plan of Jiangsu Province (No. SWYY-127), the Innovative and Entrepreneurial Talents of Jiangsu Province, the Program for High-Level Entrepreneurial and Innovative Talents Introduction of Jiangsu Province, Natural Science Foundation of Guangdong Province, Guangdong High-level Personnel of Special Support Program and Yangfan Plan of Talents Recruitment Grant, Taihu Lake Talent Plan, and Fundamental Research Funds for the Central Universities (JUSRP51712B and JUSRP1901XNC), Chinese Postdoctoral Science Foundation (No.2018M642156) and Jiangsu Planned Projects for Postdoctoral Research Funds (No.2018K040A), Youth fundation of Jiangsu Natural Science Foundation (No. BK20190599).

## Contributors

YL and JC designed the project and drafted the manuscript. JR, PH, EL and JC built up the collaboration network. JW, XP, QY and PH collected patients and contributed to the diagnosis of patients. ZX and JP extracted genomic DNA and performed exome data analysis together with JC. RG, SL, JZ performed Sanger sequencing. YL and CR contributed to Mini-gene assay and WB. JR, PH and JC reviewed study design. All of the authors revised and approved the final manuscript.

## Patient consent

Obtained.

