## Supplementary figure legend and tables for "Novel missense mutation E585K in retinitis pigmentosa leads to compromised *RPGR* splicing diversity"

**Table S1. Sequences of primers.**

| No. | Primer | Gene symbol | Exon | Primer Sequence | Used in |
| --- | --- | --- | --- | --- | --- |
| Primer pairs used for Sanger validation | | | | | |
| 1 | E585K-F | RPGR | 14 | CTTCTGCTTCTGATTCCTTCTGAC | Patient1 |
| 2 | E585K-F | RPGR | 14 | ACTGACGCAGGATACAGCTCTTAC |  |
| 3 | ORF15-F | RPGR | ORF15 | GGGATGGAACCTGTGAGGAAGGTAG | Patient2 |
| 4 | ORF15-R | RPGR | ORF15 | AATGAGTGCCCGTTATATGCAAGG |  |
| Primer pairs used for splicing assay | | | | | |
| 5 | RPGR-miniF | RPGR |  | CCGGTCGCCACCATGAAACAACAAACAATTG | Mini-gene construction |
| 6 | RPGR-miniR | RPGR |  | CCTCGCCCTTGCTCACCTCTGCTTTGTCTGTAAG |  |
| 7 | EGFP-miniF | EGFP backbone |  | TACAGACAAAGCAGAGGTGAGCAAGGGCGAGG | Mini-gene construction |
| 8 | EGFP-miniR | EGFP backbone |  | CAATTGTTTGTTGTTTCATGGTGGCGACCGG |  |
| 9 | RPGR-F | RPGR |  | ACTGACGCAGGATACAGCTCTTAC | In vitro transcription |
| 10 | RPGR-R | RPGR |  | CTGCTTTGTCTGTAAGGTCATCTG | In vitro transcription |
| 11 | Vec-R | RPGR+EGFP |  | AGTCGTGCTGCTTCATGTGGTC |  |

**Table S2. Summary of original exome sequencing data of the two RP patients in the current study.**

| **Data** | **Patient1** | **Patient2** |
| --- | --- | --- |
| Number of raw reads (M) | 146.0 | 86.3 |
| Average read length (bp) | 150 | 150 |
| Raw data yield (Gb) | 21.9 | 12.9 |
| Number of reads mapped to the genome (M) | 140.9 | 83.2 |
| Fraction of uniquely mapped bases on target (%) | 96.5% | 96.4% |
| Data mapped to target region (Gb) | 21.1 | 12.4 |
| Mean depth of target region (fold) | 163.7 | 102.4 |
| Coverage of target region (%) | 98.5% | 98.7% |
| target region with more than 10X (%) | 97.8% | 98.2% |

**Table S3. Summary of detected variants in the two RP patients.**

| Variants | **Patient1** | **Patient2** |
| --- | --- | --- |
| Number of SNPs | 108475 | 80092 |
| Number of coding SNPs | 21270 | 21163 |
| Number of synonymous SNPs | 10545 | 10526 |
| Number of nonsynonymous SNPs | 9285 | 9321 |
| Number of coding Indels | 753 | 692 |

**Table S4. Prediction of the presumed missense mutation’s effect on splicing of Exon14 with HSF.**

| **cDNA position** | **Splice site type** | **Motif** | **New splice site** | **Wild type** | **Mutant** | **Variation (%)** |
| --- | --- | --- | --- | --- | --- | --- |
| **1749** | **Donor** | GAAGTAGgt | GAAgtaagt | **79.32** | **87.66** | **+10.51** |
| **1753** | **Donor** | TAGgtaatg | TAAgtaatg | **83.77** | **73.19** | **WT site broken**  **-12.63** |

| **cDNA position** | **Linked SR protein** | **Reference Motif**  **(value 0-100)** | **Linked SR protein** | **Mutant Motif**  **(value 0-100)** | **Variation (%)** |
| --- | --- | --- | --- | --- | --- |
| **1753** |  |  | **SRp55** | **TAAgta (76.10)** | **New site** |

**Supplementary Figure Legend**

**Figure S1.** Pedigree information of the XLRP family and clinical examination. (A). Pedigree of the XLRP family in the current study. The proband (Patient 2) is indicated with arrow. (B) Fundus examination of Patient 2 shows no sign of bone-spicule pigmentation, pointing to RP sine pigmento in the patient. (C) ERG shows bilateral unrecordable responses in the patient.

**Figure S2. RNA-seq data support the existence of the intrinsic alternative splice site and Exon14_S_ in multiple cell types.** With publicly available ENCODE RNA-seq data visualized in the UCSC Genome Browser, several cell types, including H1-hESC, K562, Hela-S3, HUVEC, were found to contained reads spanning the junction of exon 14-15 lacking the last 4 bp of exon14, which further supported the existence of the intrinsic alternative splice donor and exon14 with 4-bp deletion.

**Figure S3. RNA-seq data support the existence of retention of intron 14 in multiple cell types.** With publicly available ENCODE RNA-seq data visualized in the UCSC Genome Browser, several cell types, including H1-hESC, K562, Hela-S3, HEPG2, HUVEC, were found to contained reads spanning intron 14 of *RPGR*, which further supported the existence of the intron retention.
