## Supplementary figures and images for "Novel missense mutation E585K in retinitis pigmentosa leads to compromised *RPGR* splicing diversity"

### Figure S1

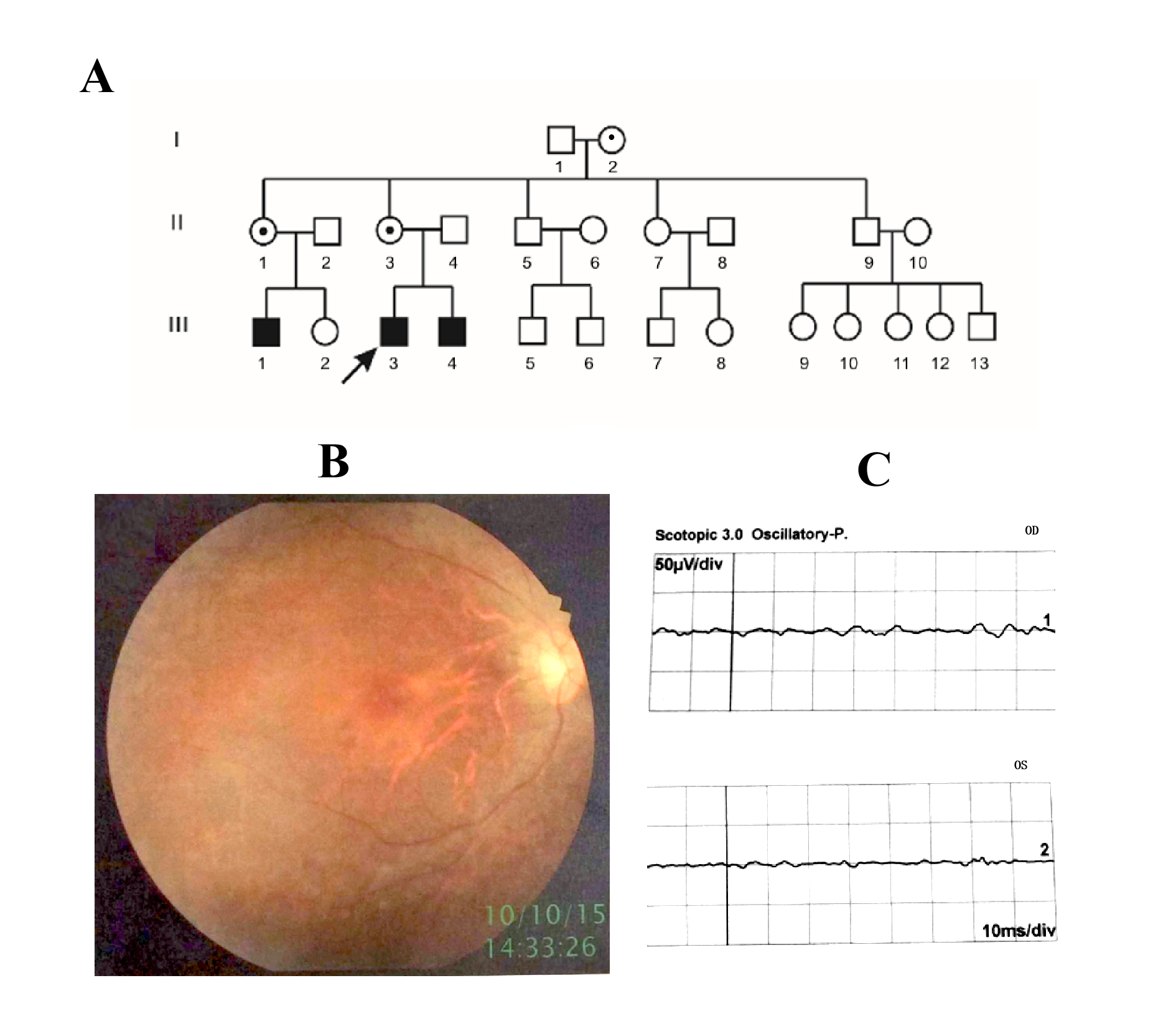

### Figure S2

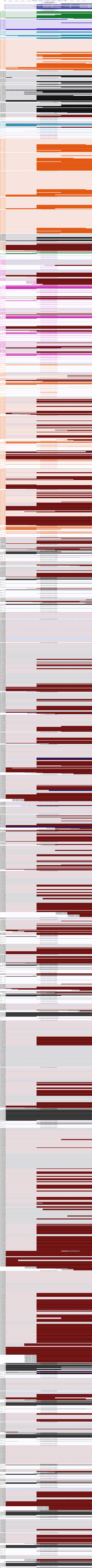

### Figure S3

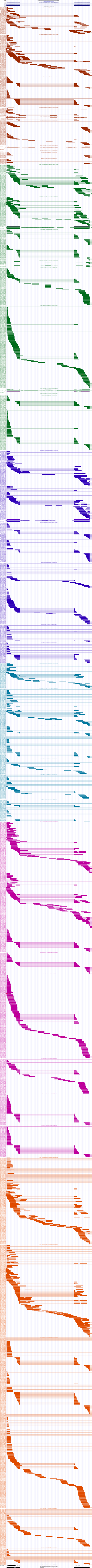
